## Supplemental figures and table for "Spatially resolved transcriptomics reveals the architecture of the tumor/microenvironment interface"

**Figure S1. Spatially resolved transcriptomics data statistics.**

**a-b.** Violin plots showing number of transcripts (UMIs) detected per Visium array spot (a) and number of unique genes detected per spot (b). **c.** Images of the H&E stained tissue sections used for SRT. **d-e.** Numbers of UMIs and genes detected per spot, plotted onto the tissue array.

**f.** Violin plot showing the number of UMIs per spot across the tissue types in the individual SRT samples. **g.** Approximation of the number of nuclei per Visium array spot (55 μm diameter) in the tumor and muscle regions.

### Figure S1

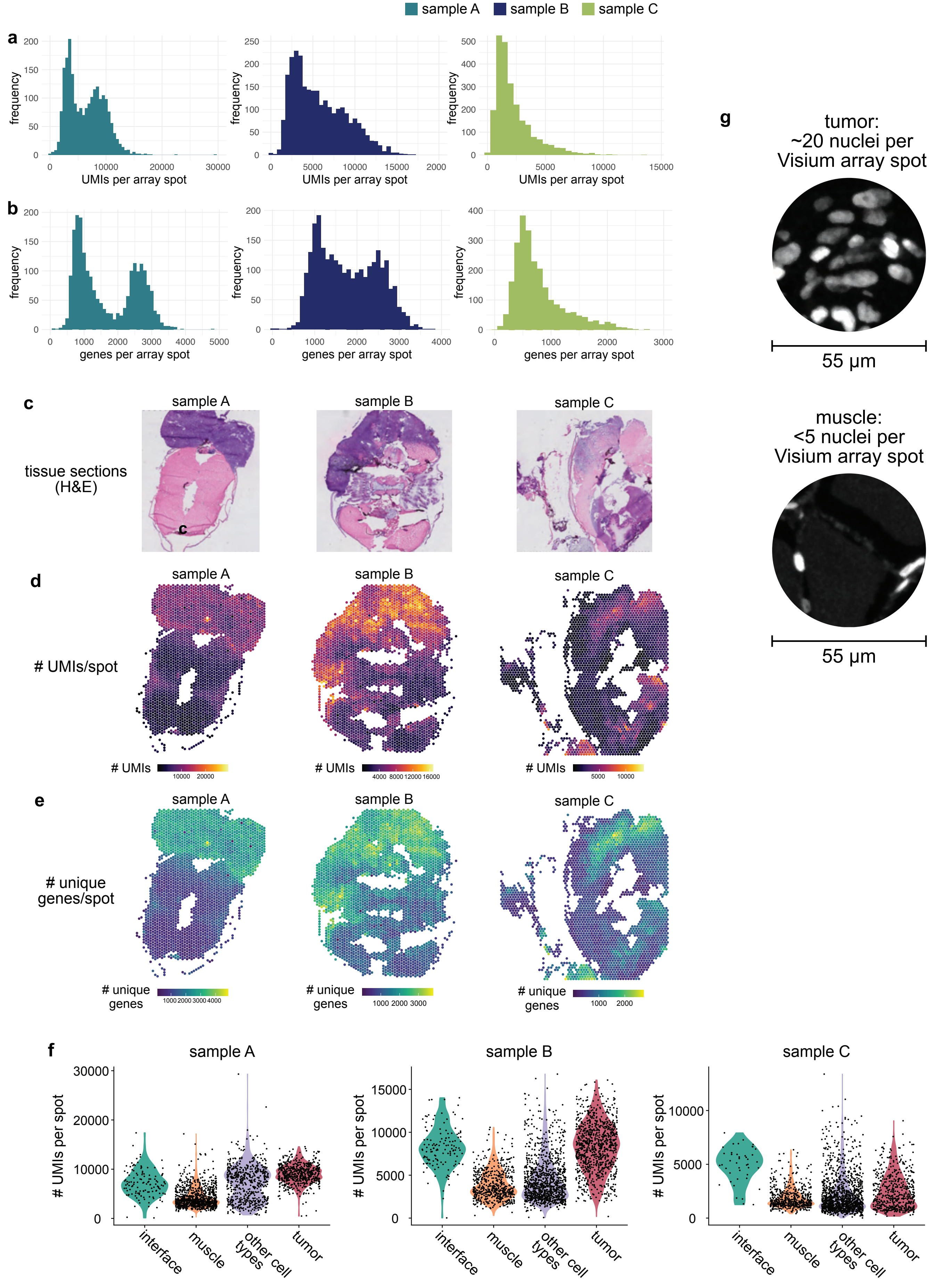

sample A
sample B
sample C
a
frequency
UMIs per array spot
b
genes per array spot
c
tissue sections (H&E)
d
### UMIs/spot
### UMIs
e
### unique genes/spot
### unique genes
f
### UMIs per spot
interface
muscle
other cell types
tumor
g
tumor: ~20 nuclei per Visium array spot
55 µm
muscle: <5 nuclei per Visium array spot
55 µm

**Figure S2. A unique interface cluster is found in each sample.**
**a.** Cluster assignments plotted onto the Visium array for individual samples. **b.** Cluster assignments labelled in UMAP space for individual samples.

**Figure S2**

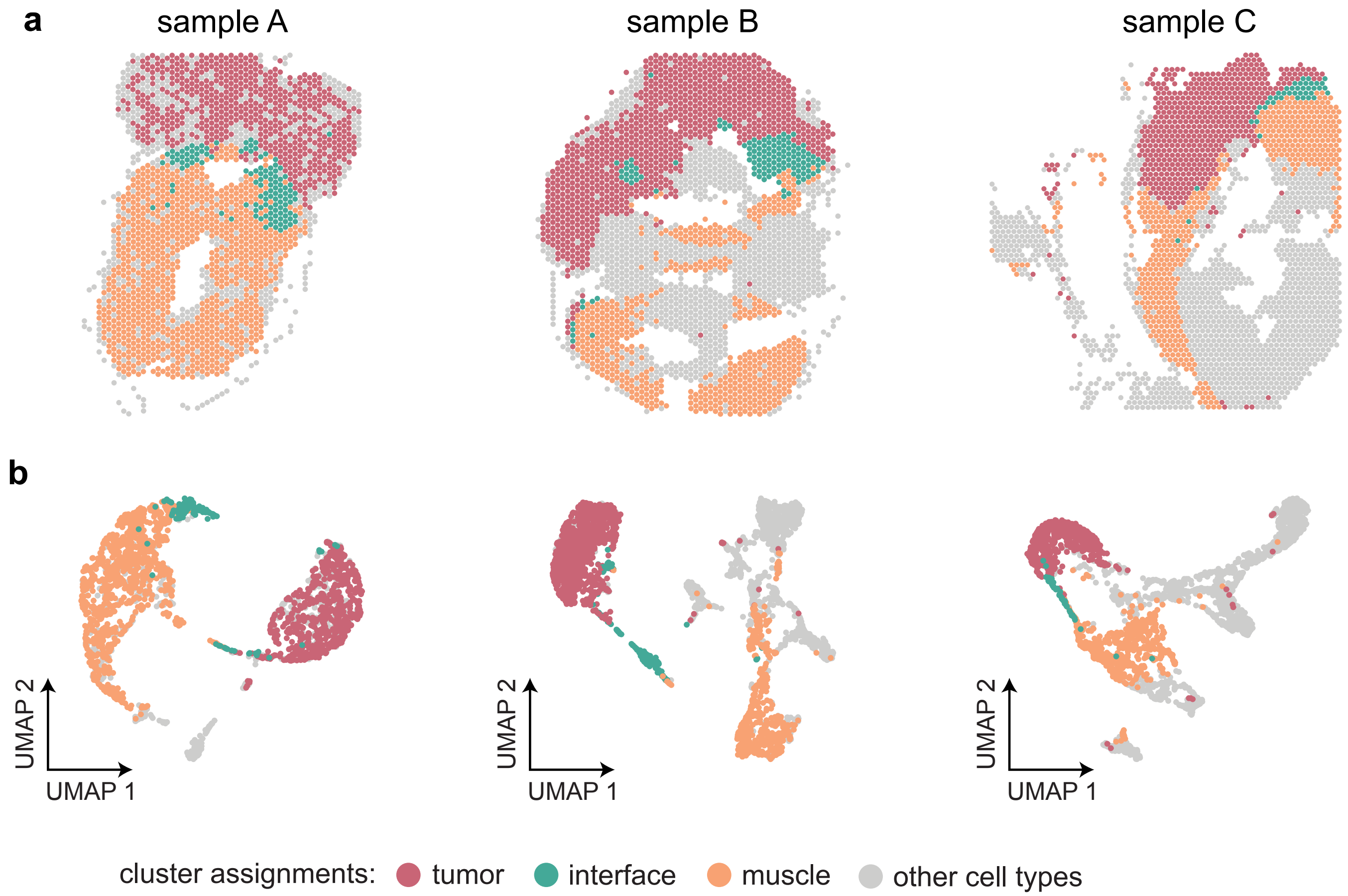
a
sample A
sample B
sample C
b
UMAP 2
UMAP 1
UMAP 2
UMAP 1
UMAP 2
UMAP 1
cluster assignments:
tumor
interface
muscle
other cell types

**Figure S3. Spatial patterning of biological pathways in the tumor and microenvironment.**
**a-b.** Average, standardized expression of annotated genes for gene ontology (GO) terms displaying spatially-coherent expression patterns in the tumor (a) and microenvironment (b) regions in each SRT sample. P-values represent the comparison between the distance between spots expressing that GO term genes and a null-distribution of distances between random spots (Wilcoxon's Rank Sum test, see Methods).

#### Figure S3

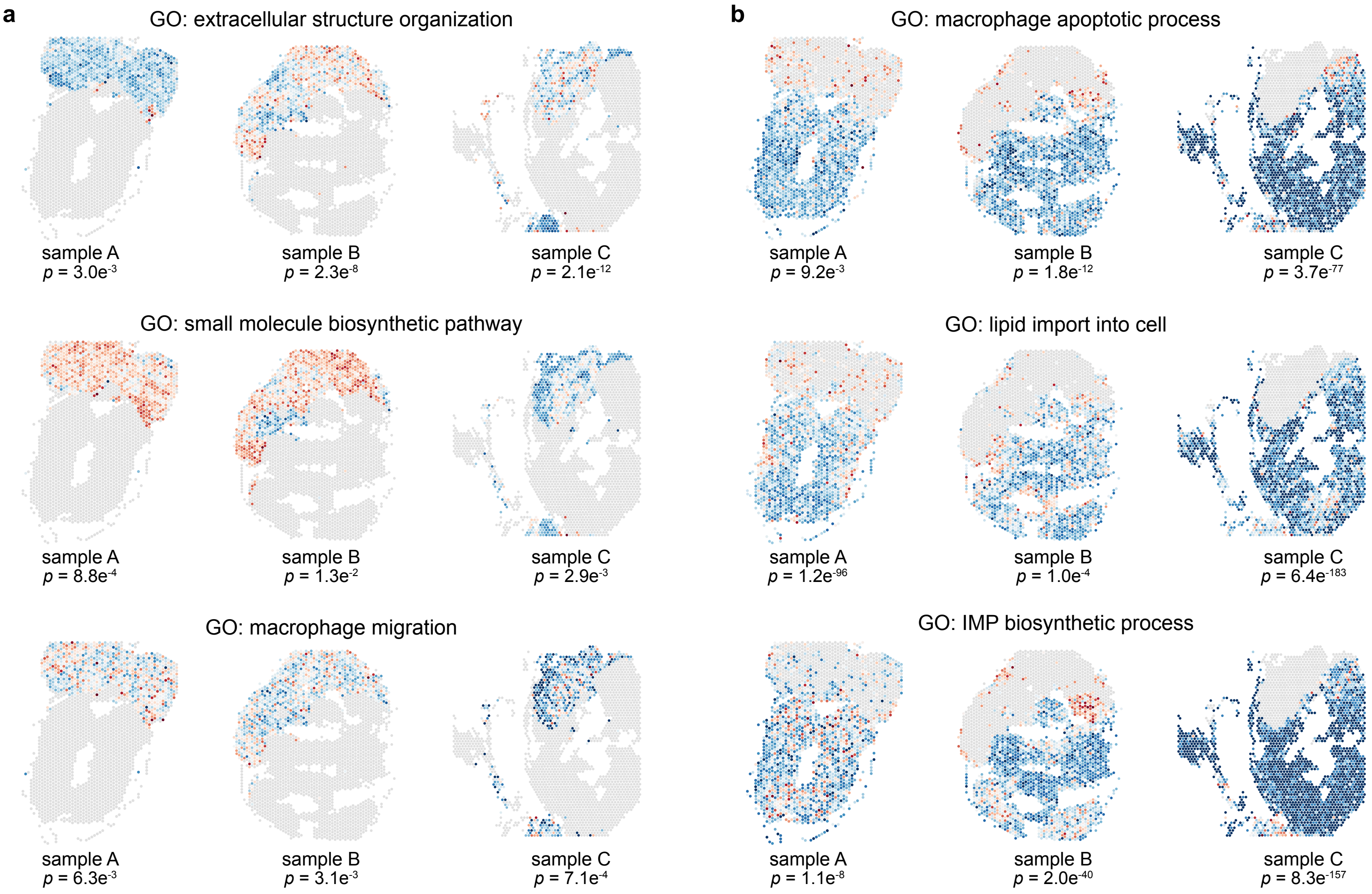

a
GO: extracellular structure organization
sample A
p = 3.0e-3
sample B
p = 2.3e-8
sample C
p = 2.1e-12
GO: small molecule biosynthetic pathway
sample A
p = 8.8e-4
sample B
p = 1.3e-2
sample C
p = 2.9e-3
GO: macrophage migration
sample A
p = 6.3e-3
sample B
p = 3.1e-3
sample C
p = 7.1e-4
b
GO: macrophage apoptotic process
sample A
p = 9.2e-3
sample B
p = 1.8e-12
sample C
p = 3.7e-77
GO: lipid import into cell
sample A
p = 1.2e-96
sample B
p = 1.0e-4
sample C
p = 6.4e-183
GO: IMP biosynthetic process
sample A
p = 1.1e-8
sample B
p = 2.0e-40
sample C
p = 8.3e-157

**Figure S4. Non-negative matrix factorization (NMF) analysis on SRT spots, and enriched GO terms for top genes in each NMF factor.**

Non-negative matrix factorization (NMF) was performed on the integrated expression matrix of the three SRT samples, with k = 11 factors. Factor scores were then projected onto spots, and the top 150 scoring genes per factor were used for GO enrichment analysis.

### Figure S4

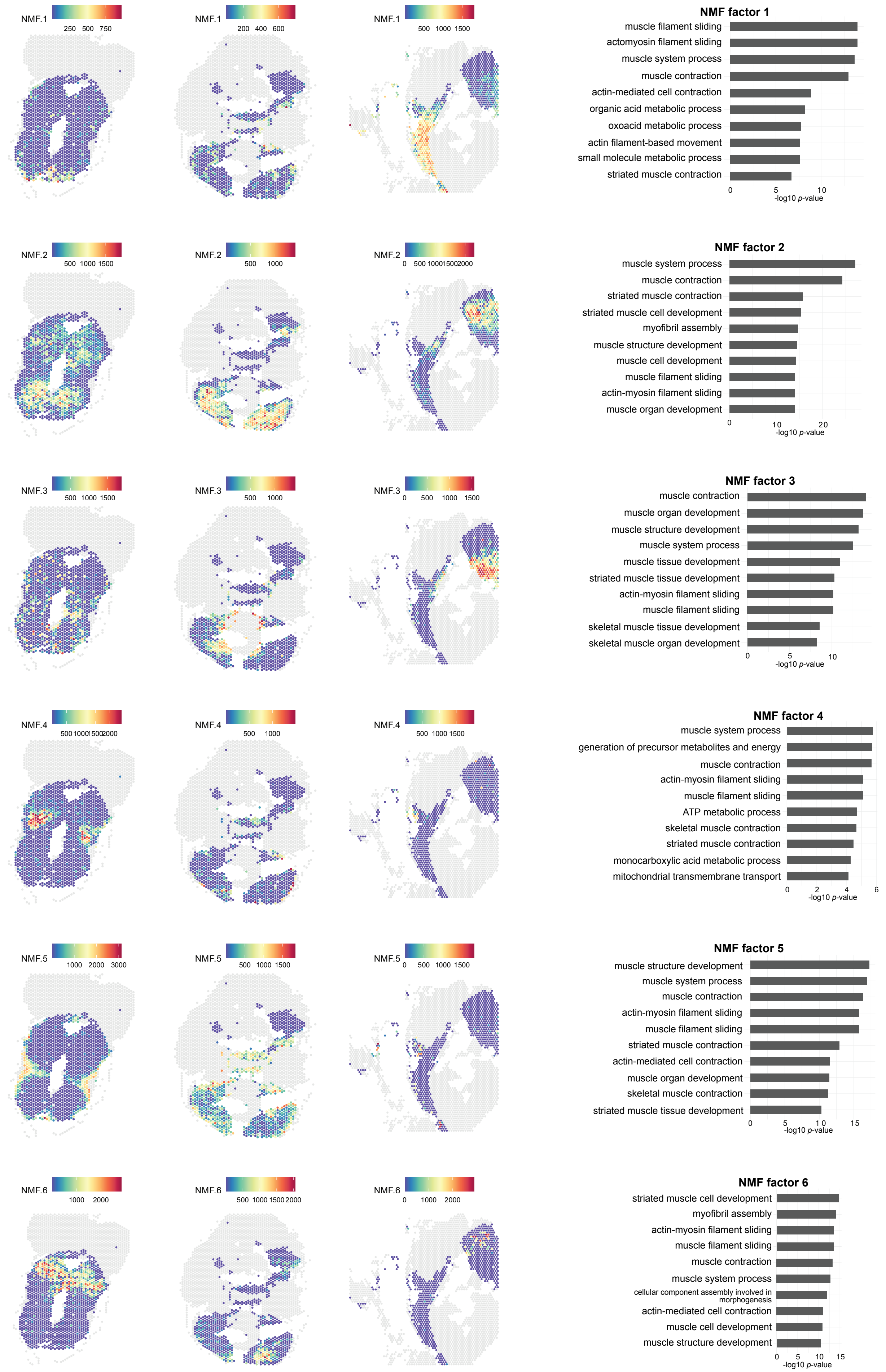
NMF.1
250 500 750
200 400 600
500 1000 1500
NMF factor 1
muscle filament sliding
actomyosin filament sliding
muscle system process
muscle contraction
actin-mediated cell contraction
organic acid metabolic process
oxoacid metabolic process
actin filament-based movement
small molecule metabolic process
striated muscle contraction
0 5 10
-log10 p-value
NMF.2
500 1000 1500
500 1000
0 500 100015002000
NMF factor 2
muscle system process
muscle contraction
striated muscle contraction
striated muscle cell development
myofibril assembly
muscle structure development
muscle cell development
muscle filament sliding
actin-myosin filament sliding
muscle organ development
0 10 20
-log10 p-value
NMF.3
500 1000 1500
500 1000
0 500 1000 1500
NMF factor 3
muscle contraction
muscle organ development
muscle structure development
muscle system process
muscle tissue development
striated muscle tissue development
actin-myosin filament sliding
muscle filament sliding
skeletal muscle tissue development
skeletal muscle organ development
0 5 10
-log10 p-value
NMF.4
500100015002000
500 1000
500 1000 1500
NMF factor 4
muscle system process
generation of precursor metabolites and energy
muscle contraction
actin-myosin filament sliding
muscle filament sliding
ATP metabolic process
skeletal muscle contraction
striated muscle contraction
monocarboxylic acid metabolic process
mitochondrial transmembrane transport
0 2 4 6
-log10 p-value
NMF.5
1000 2000 3000
500 1000 1500
0 500 1000 1500
NMF factor 5
muscle structure development
muscle system process
muscle contraction
actin-myosin filament sliding
muscle filament sliding
striated muscle contraction
actin-mediated cell contraction
muscle organ development
skeletal muscle contraction
striated muscle tissue development
0 5 10 15
-log10 p-value
NMF.6
1000 2000
500 1000 1500 2000
0 1000 2000
NMF factor 6
striated muscle cell development
myofibril assembly
actin-myosin filament sliding
muscle filament sliding
muscle contraction
muscle system process
cellular component assembly involved in morphogenesis
actin-mediated cell contraction
muscle cell development
muscle structure development
0 5 10 15
-log10 p-value

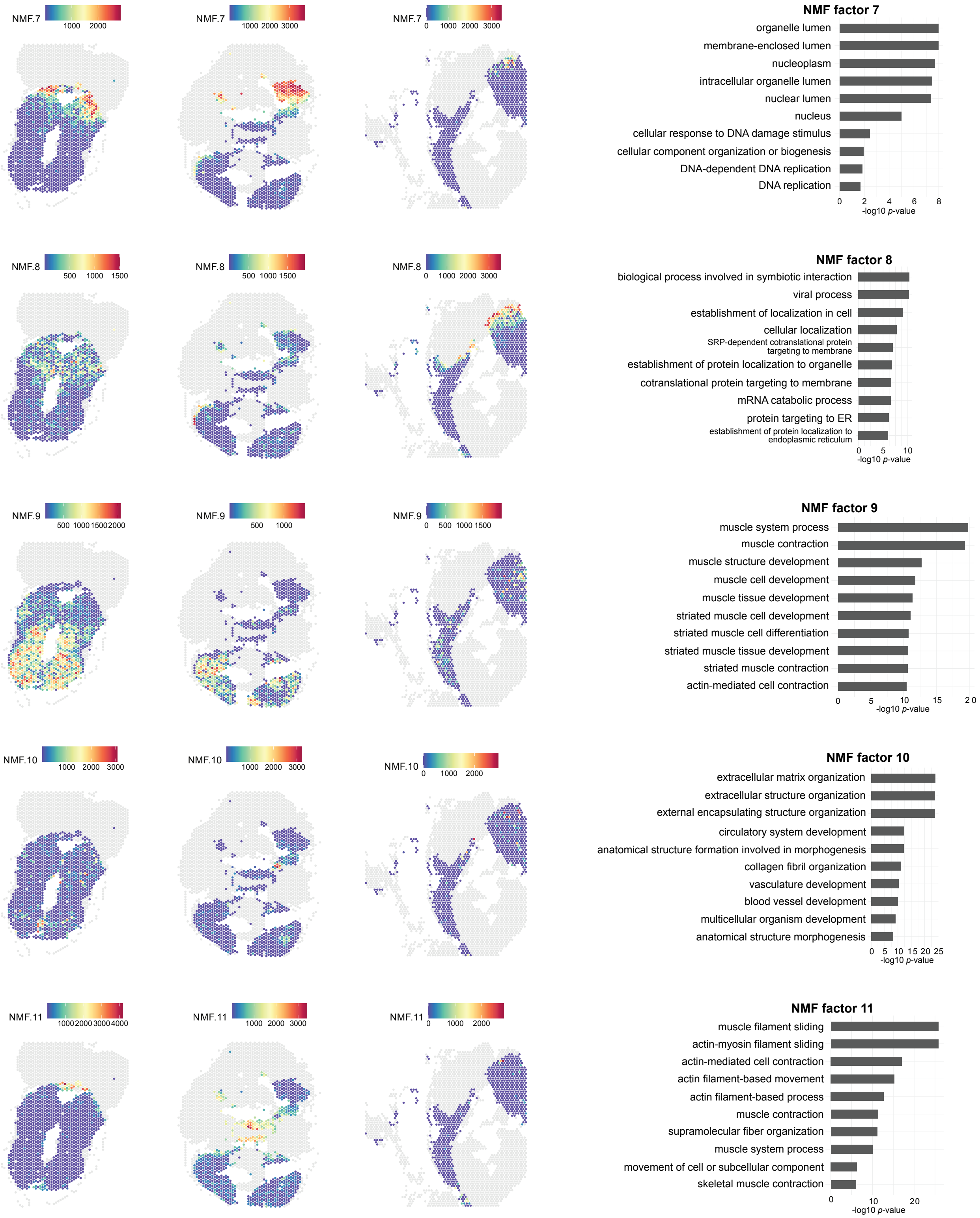
NMF.7
1000 2000
1000 2000 3000
0 1000 2000 3000
NMF factor 7
organelle lumen
membrane-enclosed lumen
nucleoplasm
intracellular organelle lumen
nuclear lumen
nucleus
cellular response to DNA damage stimulus
cellular component organization or biogenesis
DNA-dependent DNA replication
DNA replication
0 2 4 6 8
-log10 p-value
NMF.8
500 1000 1500
0 1000 2000 3000
NMF factor 8
biological process involved in symbiotic interaction
viral process
establishment of localization in cell
cellular localization
SRP-dependent cotranslational protein targeting to membrane
establishment of protein localization to organelle
cotranslational protein targeting to membrane
mRNA catabolic process
protein targeting to ER
establishment of protein localization to endoplasmic reticulum
0 5 10
-log10 p-value
NMF.9
500 1000 1500 2000
500 1000
0 500 1000 1500
NMF factor 9
muscle system process
muscle contraction
muscle structure development
muscle cell development
muscle tissue development
striated muscle cell development
striated muscle cell differentiation
striated muscle tissue development
striated muscle contraction
actin-mediated cell contraction
0 5 10 15 20
-log10 p-value
NMF.10
1000 2000 3000
0 1000 2000
NMF factor 10
extracellular matrix organization
extracellular structure organization
external encapsulating structure organization
circulatory system development
anatomical structure formation involved in morphogenesis
collagen fibril organization
vasculature development
blood vessel development
multicellular organism development
anatomical structure morphogenesis
0 5 10 15 20 25
-log10 p-value
NMF.11
1000 2000 3000 4000
1000 2000 3000
0 1000 2000
NMF factor 11
muscle filament sliding
actin-myosin filament sliding
actin-mediated cell contraction
actin filament-based movement
actin filament-based process
muscle contraction
supramolecular fiber organization
muscle system process
movement of cell or subcellular component
skeletal muscle contraction
0 10 20
-log10 p-value

**Figure S5. single-cell RNA-seq data statistics. a,d.** Histograms showing the number of UMIs (left) and genes (right) per cell for each scRNA-seq reaction. **b,e.** Violin plots showing the number of UMIs (left) and genes (right) per cell for the different clusters in each scRNA-seq reaction. **c,f.** Cluster assignments for each scRNA-seq reaction plotted in UMAP space. **g.** Predicted possible doublets within the integrated scRNA-seq dataset.

### Figure S5

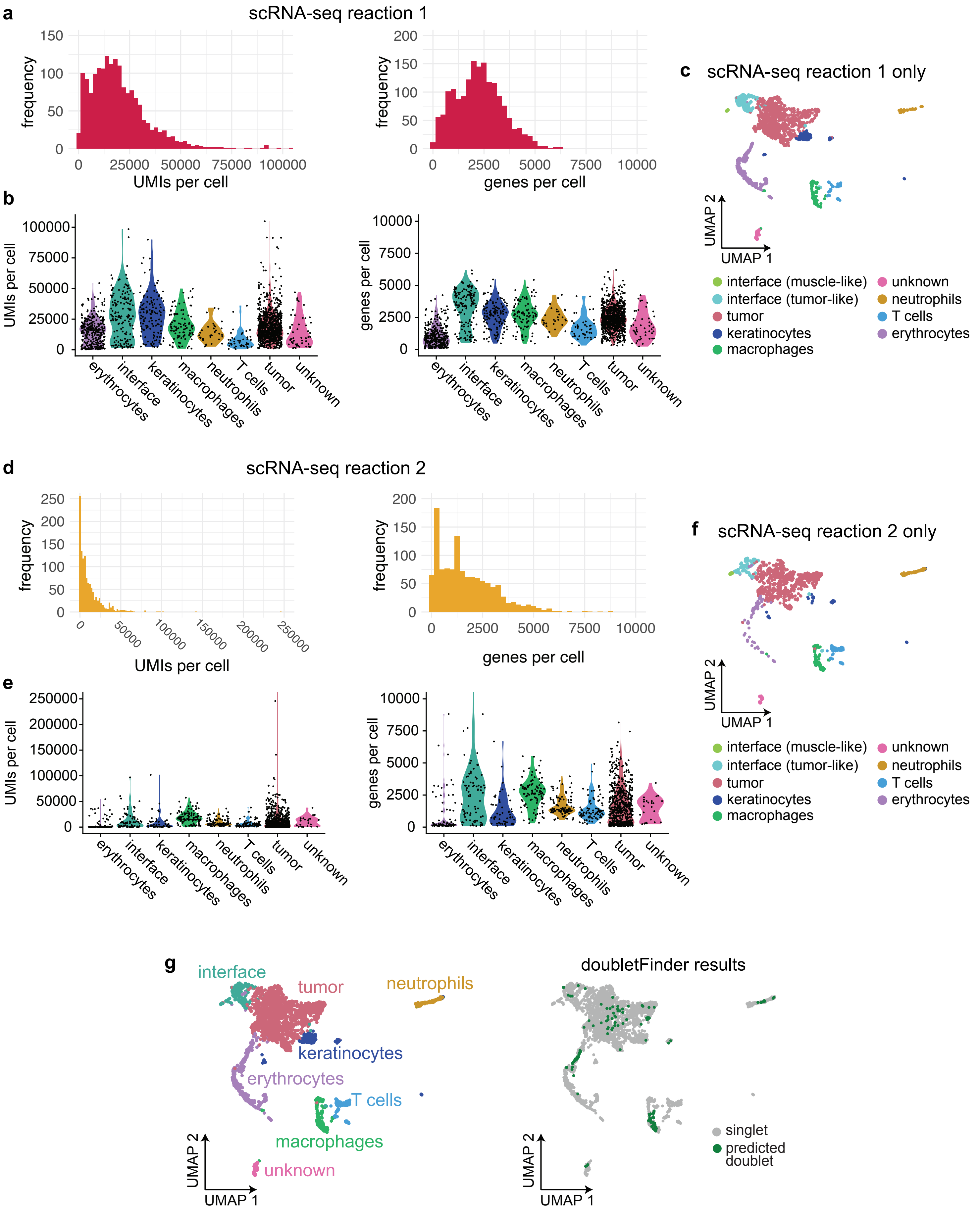

a
scRNA-seq reaction 1
frequency
UMIs per cell
genes per cell
b
UMIs per cell
genes per cell
erythrocytes
interface
keratinocytes
macrophages
neutrophils
T cells
tumor
unknown
c
scRNA-seq reaction 1 only
UMAP 1
UMAP 2
interface (muscle-like)
interface (tumor-like)
tumor
keratinocytes
macrophages
unknown
neutrophils
T cells
erythrocytes
d
scRNA-seq reaction 2
frequency
UMIs per cell
genes per cell
e
UMIs per cell
genes per cell
f
scRNA-seq reaction 2 only
g
interface
tumor
neutrophils
keratinocytes
erythrocytes
T cells
macrophages
unknown
doubletFinder results
singlet
predicted doublet

**Figure S6. single-nucleus RNA-seq data statistics. a.** Histograms showing the number of UMIs (left) and genes (right) per nucleus. **b.** Violin plots showing the number of UMIs (left) and genes (right) per nucleus across the clusters in the snRNA-seq dataset. **c-d.** Mean expression per cell/nucleus for all common genes in the scRNA-seq and snRNA-seq dataset (15,022 genes) (c) and for the fish orthologs of the published SYSCILIA genes (320 genes).

#### Figure S6

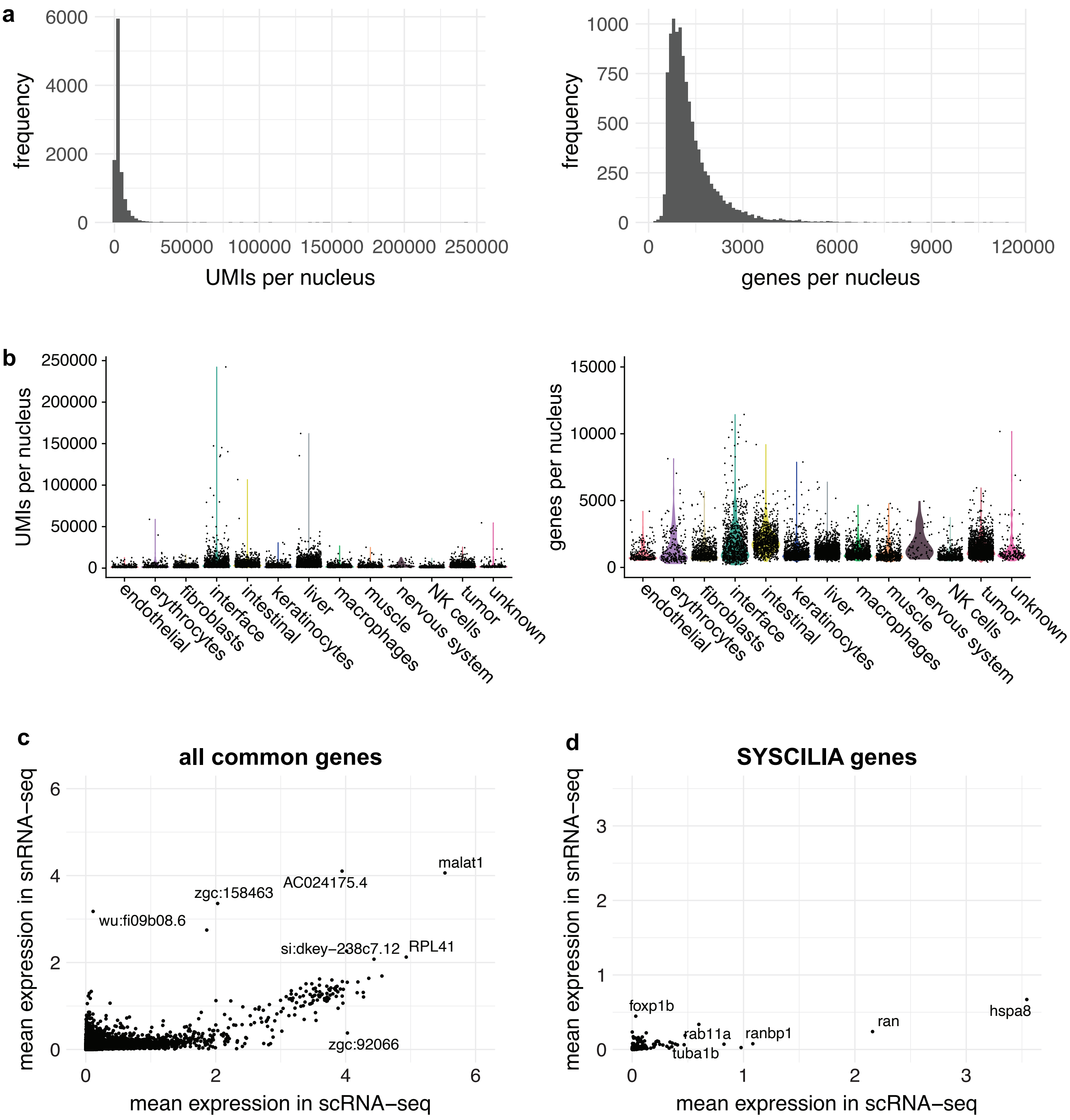
a
frequency
UMIs per nucleus
0
2000
4000
6000
0
50000
100000
150000
200000
250000
frequency
genes per nucleus
0
250
500
750
1000
0
3000
6000
9000
120000
b
UMIs per nucleus
genes per nucleus
endothelial
erythrocytes
fibroblasts
interface
intestinal
keratinocytes
liver
macrophages
muscle
nervous system
NK cells
tumor
unknown
c
all common genes
mean expression in snRNA−seq
mean expression in scRNA−seq
malat1
AC024175.4
zgc:158463
wu:fi09b08.6
si:dkey−238c7.12
RPL41
zgc:92066
d
SYSCILIA genes
foxp1b
rab11a
tuba1b
ranbp1
ran
hspa8

**Figure S7. Deconvolution of the SRT interface region using multimodal intersection analysis (MIA). a-b.** MIA maps for all snRNA-seq clusters compared to all SRT clusters (a) or only the SRT interface cluster (b). *P*-values were calculated using the hypergeometric test.

#### Figure S7

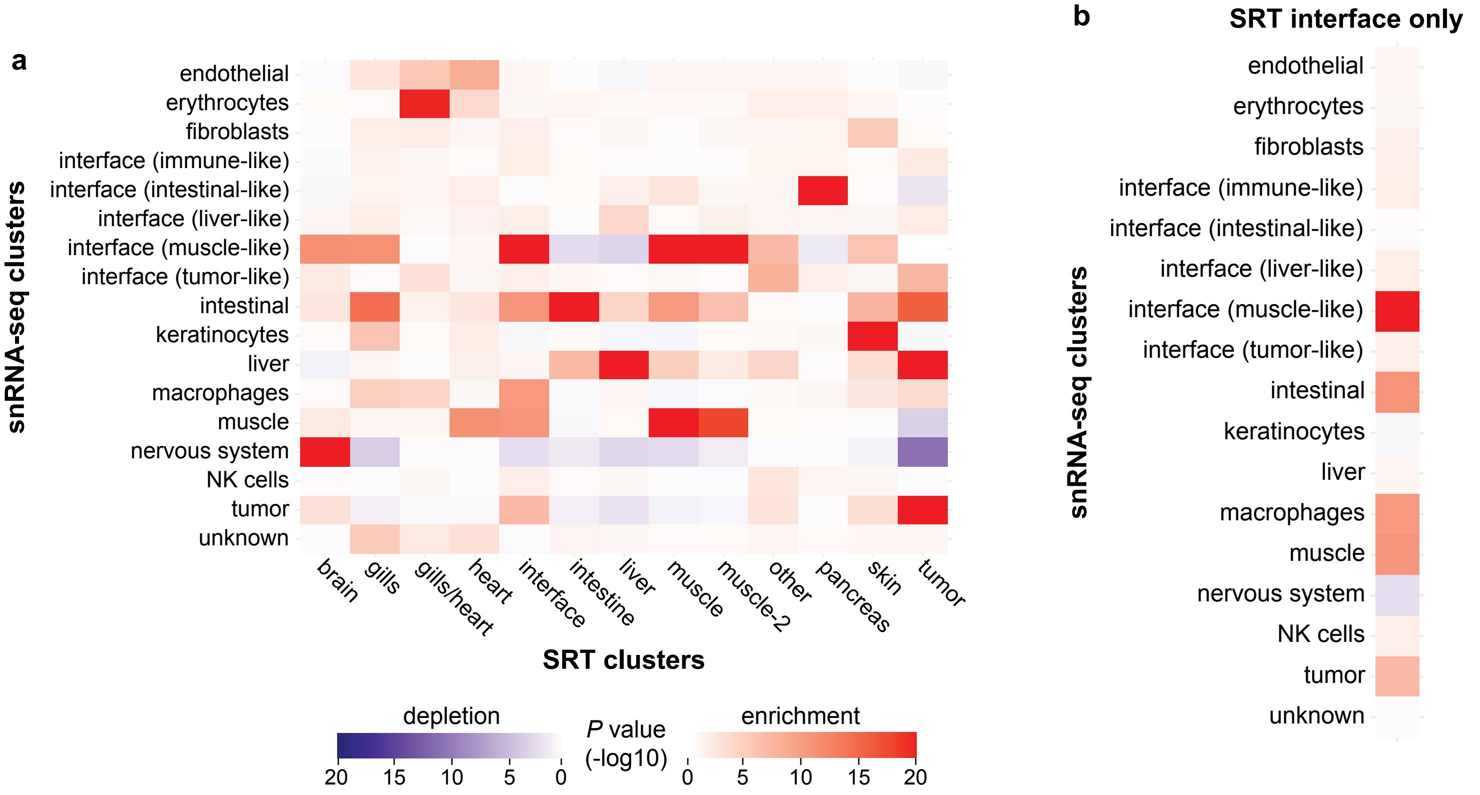
a
endothelial
erythrocytes
fibroblasts
interface (immune-like)
interface (intestinal-like)
interface (liver-like)
interface (muscle-like)
interface (tumor-like)
intestinal
keratinocytes
liver
macrophages
muscle
nervous system
NK cells
tumor
unknown
snRNA-seq clusters
brain
gills
gills/heart
heart
interface
intestine
liver
muscle
muscle-2
other
pancreas
skin
tumor
SRT clusters
depletion
20 15 10 5 0
P value (-log10)
enrichment
0 5 10 15 20
b
SRT interface only
endothelial
erythrocytes
fibroblasts
interface (immune-like)
interface (intestinal-like)
interface (liver-like)
interface (muscle-like)
interface (tumor-like)
intestinal
keratinocytes
liver
macrophages
muscle
nervous system
NK cells
tumor
unknown
snRNA-seq clusters

**Figure S8. Computational modeling of interface cell interactions with NicheNet. a.** Predicted ligands expressed by interface cells, ranked by predicted ligand activity (Pearson correlation coefficient). **b.** Predicted receptors expressed by non-interface cells for the predicted ligands in **a. c.** Predicted target genes for the ligands in **a**. **d-e.** Normalized expression of the zebrafish orthologs of *HMGB2* (*hmgb2a* and *hmgb2b*) and *CDH1* (*cdh1*) in the snRNA-seq, scRNA-seq and SRT clusters.

### Figure S8

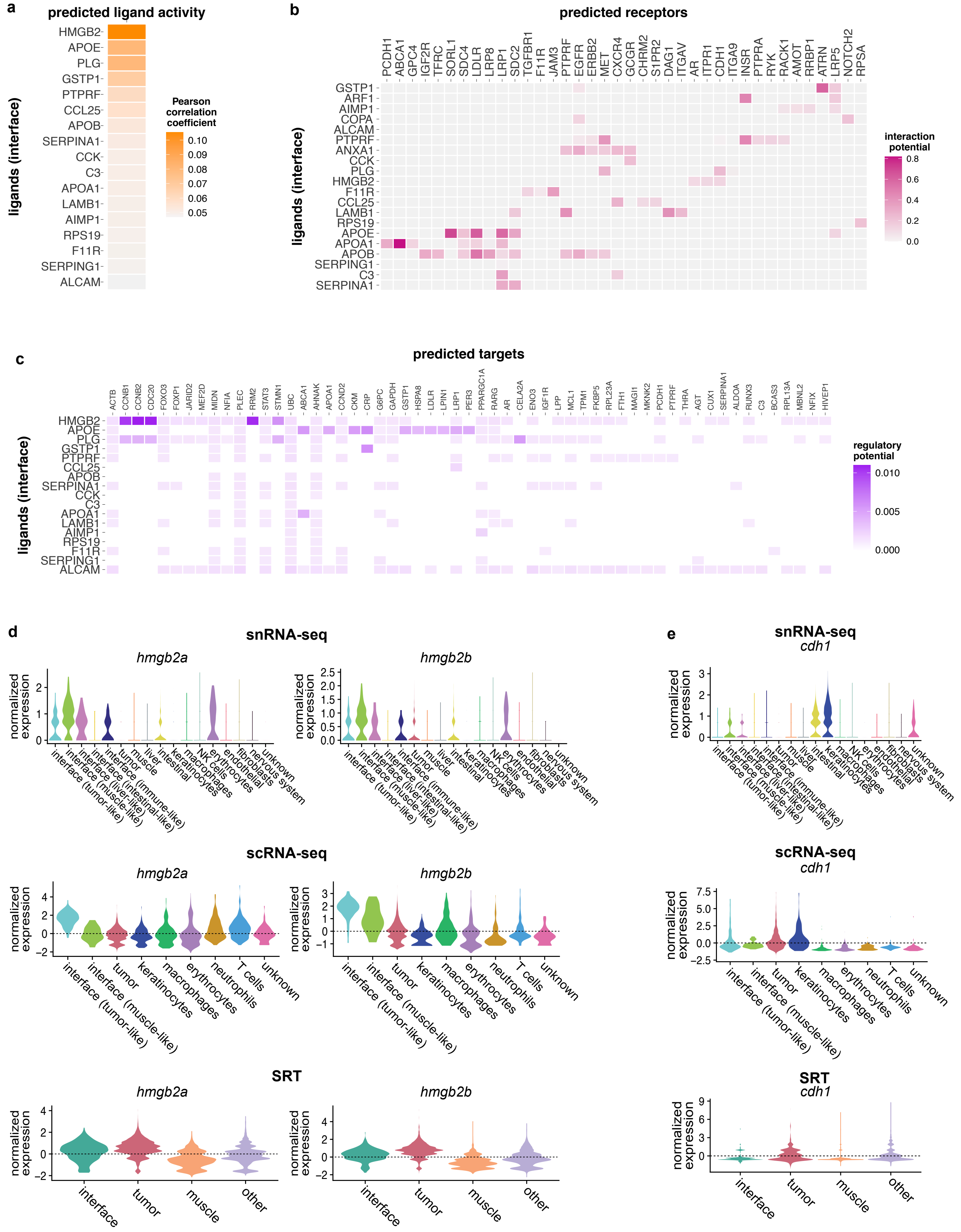
a
predicted ligand activity
ligands (interface)
HMGB2
APOE
PLG
GSTP1
PTPRF
CCL25
APOB
SERPINA1
CCK
C3
APOA1
LAMB1
AIMP1
RPS19
F11R
SERPING1
ALCAM
Pearson correlation coefficient
0.10
0.09
0.08
0.07
0.06
0.05
b
predicted receptors
PCDH1
ABCA1
GPC4
IGF2R
TFRC
SORL1
SDC4
LDLR
LRP8
LRP1
SDC2
TGFBR1
F11R
JAM3
PTPRF
EGFR
ERBB2
MET
CXCR4
GCGR
CHRM2
S1PR2
DAG1
ITGAV
AR
ITPR1
CDH1
ITGA9
INSR
PTPRA
RYK
RACK1
AMOT
RRBP1
ATRN
LRP5
NOTCH2
RPSA
GSTP1
ARF1
AIMP1
COPA
ALCAM
PTPRF
ANXA1
CCK
PLG
HMGB2
F11R
CCL25
LAMB1
RPS19
APOE
APOA1
APOB
SERPING1
C3
SERPINA1
interaction potential
0.8
0.6
0.4
0.2
0.0
c
predicted targets
ACTB
CCNB1
CCNB2
CDC20
FOXO3
FOXP1
JARID2
MEF2D
MIDN
NFIA
PLEC
RRM2
STAT3
STMN1
UBC
ABCA1
AHNAK
APOA1
CCND2
CKM
CRP
G6PC
GAPDH
GSTP1
HSPA8
LDLR
LPIN1
LRP1
PER3
PPARGC1A
RARG
AR
CELA2A
ENO3
IGF1R
LPP
MCL1
TPM1
FKBP5
RPL23A
FTH1
MAGI1
MKNK2
PCDH1
PTPRF
THRA
AGT
CUX1
SERPINA1
ALDOA
RUNX3
C3
BCAS3
RPL13A
MBNL2
NFIX
HIVEP1
regulatory potential
0.010
0.005
0.000
d
snRNA-seq
hmgb2a
hmgb2b
normalized expression
interface (tumor-like)
interface (muscle-like)
interface (liver-like)
interface (intestinal-like)
interface (immune-like)
tumor
muscle
liver
intestinal
keratinocytes
macrophages
NK cells
erythrocytes
endothelial
fibroblasts
nervous system
unknown
scRNA-seq
interface (tumor-like)
interface (muscle-like)
tumor
keratinocytes
macrophages
erythrocytes
neutrophils
T cells
unknown
SRT
interface
tumor
muscle
other
e
snRNA-seq
cdh1
scRNA-seq
cdh1
SRT
cdh1

**Figure S9. Acetylated tubulin staining within the tumor, muscle, and tumor-muscle interface regions.** Immunofluorescence images of tumor (green), Hoescht (blue) and acetylated tubulin (magenta) intensity at the tumor-muscle interface (top), center of tumor (middle) and distant muscle (bottom). Scale bars, 100 μm (40X magnification).

#### Figure S9

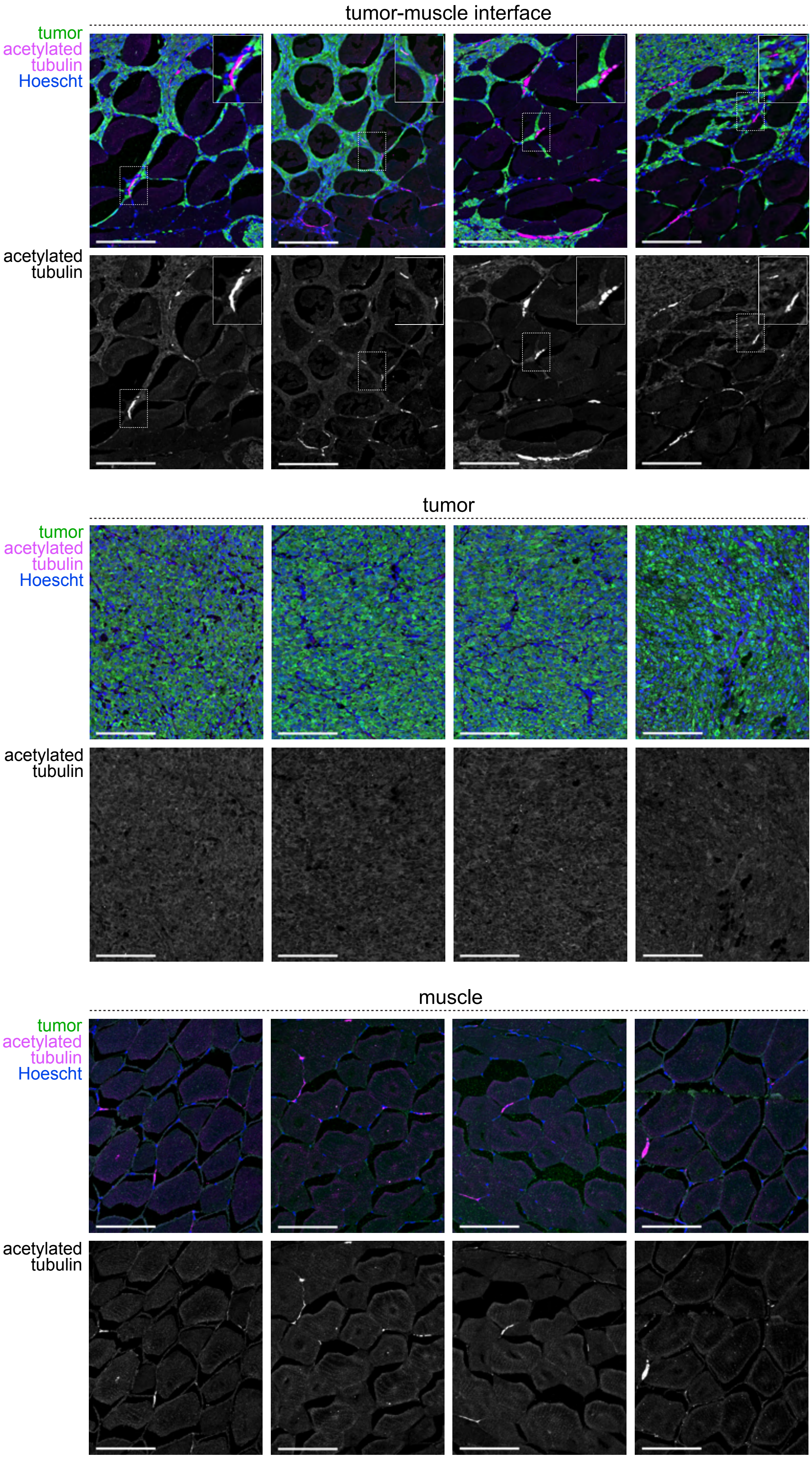
tumor-muscle interface
tumor
acetylated
tubulin
Hoescht
acetylated
tubulin
tumor
tumor
acetylated
tubulin
Hoescht
acetylated
tubulin
muscle
tumor
acetylated
tubulin
Hoescht
acetylated
tubulin

**Figure S10. An interface-like cell state is found in human melanoma. a-c.** UMAP projection of human melanoma scRNA-seq data from Tirosh et al., 2016. Cell types (original annotations, a), interface marker gene expression score (b), and interface classification (c) are indicated. The cutoff for classification as an interface-like cell state is indicated in (b). **d-e.** Expression of human cilia (d) and ETS (e) genes across the cell types/states in human melanoma scRNA-seq. The expression score represents the standardized mean expression of all indicated genes per cell.

### Figure S10

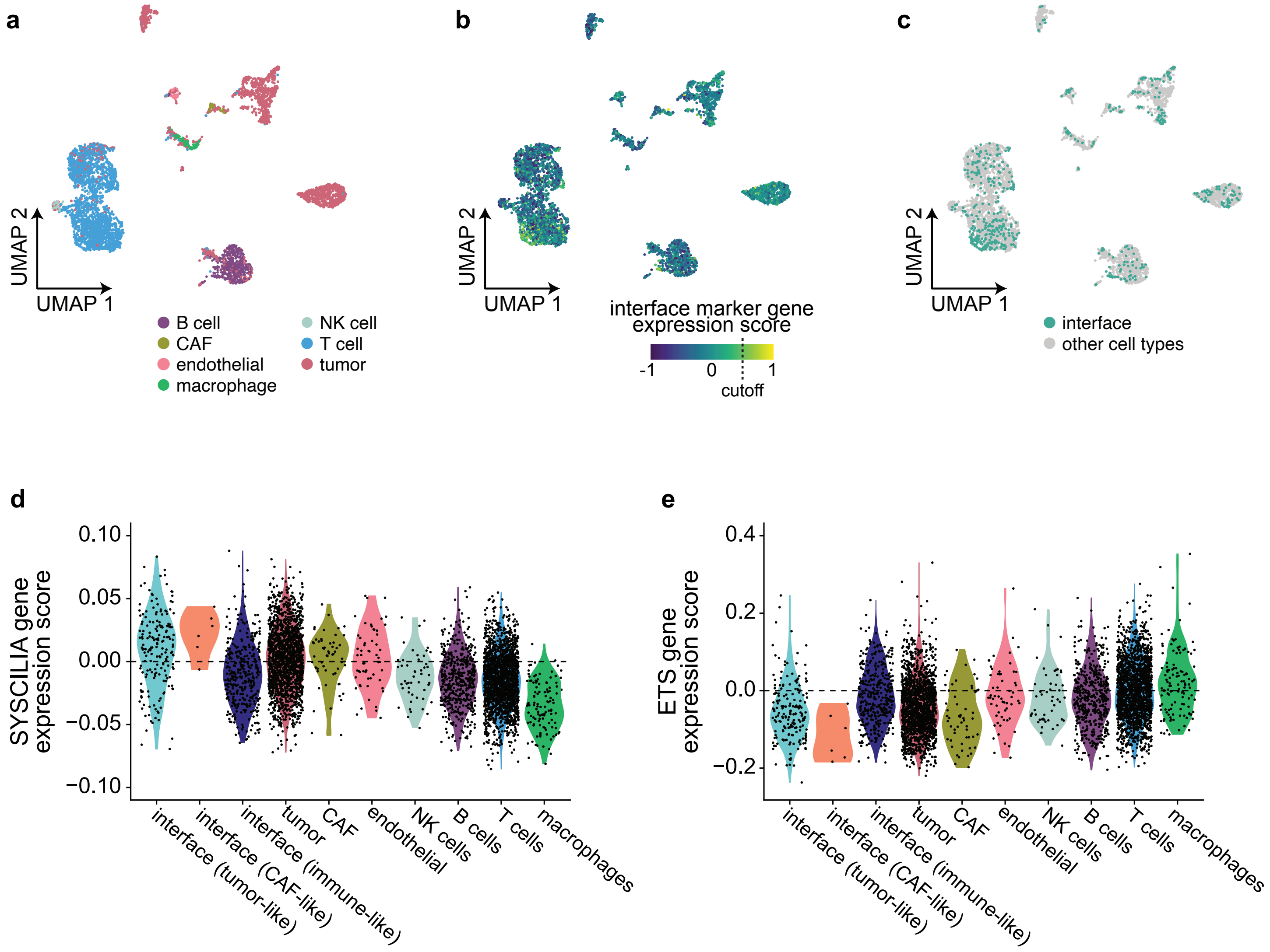

a
b
c
UMAP 2
UMAP 1
B cell
CAF
endothelial
macrophage
NK cell
T cell
tumor
interface marker gene
expression score
-1
0
1
cutoff
interface
other cell types
d
e
SYSCILIA gene
expression score
0.10
0.05
0.00
−0.05
−0.10
ETS gene
expression score
0.4
0.2
0.0
−0.2
interface (tumor-like)
interface (CAF-like)
interface (immune-like)
tumor
CAF
endothelial
NK cells
B cells
T cells
macrophages

**Table S1. Activator/repressor status for the human ETS genes.** Activator/repressor status and fish orthologs were obtained from Uniprot and ZFIN.

#### Table S1

| Human gene | Fish ortholog(s) | Activator or repressor? |
|---|---|---|
| *ERF* | *erf, erfl1, erfl3* | Repressor |
| *ELK1* | *elk1* | Activator |
| *ELK3* | *elk3* | Either |
| *ELK4* | *elk4* | Either |
| *ERG* | *erg* | Unknown |
| *FLI1* | *fli1, fli1rs* | Activator |
| *FEV* | *fev* | Repressor |
| *ETS1* | *ets1* | Activator |
| *ETS2* | *ets2* | Activator |
| *ETV2* | *etsrp* | Activator |
| *GABPA* | *gabpa* | Activator |
| *ETV5* | *etv5a, etv5b* | Activator |
| *ETV1* | *etv1* | Activator |
| *ETV4* | *etv4* | Activator |
| *SPDEF* | *spdef* | Activator |
| *EHF* | *ehf* | Either |
| *ELF3* | *elf3* | Either |
| *ELF4* | *elf1* | Activator |
| *ELF2* | *elf2a, elf2b* | Either |
| *ETV6* | *etv6* | Repressor |
| *ETV7* | *etv7* | Repressor |
| *SPI1* | *spi1b* | Activator |
| *SPIC* | *spic, spicl1* | Activator |
